# Genomic Insights Guiding Personalized First-Line Immunotherapy Response in Lung and Bladder Tumors

**DOI:** 10.1101/2024.10.25.620156

**Authors:** Jenifer Brea-Iglesias, María Gallardo-Gómez, Ana Oitabén, Martin E. Lázaro-Quintela, Luis León, Joao M. Alves, Laura Juaneda-Magdalena, Carme García-Benito, Ihab Abdulkader, Laura Muinelo, Jesús M Paramio, Mónica Martínez-Fernández

## Abstract

Immune checkpoint inhibitors (ICIs) have revolutionized cancer treatment, particularly in advanced non-small cell lung cancer (NSCLC) and muscle-invasive bladder cancer (MIBC). However, identifying reliable predictive biomarkers for ICI response remains a significant challenge. In this study, we analyzed real-world cohorts of advanced NSCLC and MIBC patients treated with ICIs as first-line therapy. Tumor samples underwent Whole Genome Sequencing (WGS) to identify specific somatic variants and assess tumor mutational burden (TMB). Additionally, mutational signature extraction and pathway enrichment analyses were performed to uncover the underlying mechanisms of ICI response. We also characterized HLA-I haplotypes and investigated LINE-1 retrotransposition. Distinct mutation patterns were identified in patients who responded to treatment, suggesting potential biomarkers for predicting ICI effectiveness. In NSCLC, tumor mutational burden (TMB) did not differ significantly between responders and non-responders, while in MIBC, higher TMB was linked to better responses. Specific mutational signatures and HLA haplotypes were associated with ICI response in both cancers. Pathway analysis showed that NSCLC responders had active inflammatory and immune pathways, while non-responders had pathways such related to FGFR3 and neural crest differentiation associated to resistance mechanisms. In MIBC, responders had alterations in DNA repair, leading to more neoantigens and a stronger ICI response. Importantly, for the first time, we found that LINE-1 activation was positively linked to ICI response, especially in MIBC. These findings reveal promising biomarkers and mechanistic insights, offering a new perspective on predicting ICI response and opening up exciting possibilities for more personalized immunotherapy strategies in NSCLC and MIBC.

## INTRODUCTION

Immune checkpoint inhibitors (ICI) are improving patient survival and enhancing available therapeutic options for different cancers. Their association with improved quality of life and longer life expectancy in advanced cancer patients, whose prior therapeutic options were limited, has positioned them as the standard first-line treatment for the management of several advanced tumors. Nevertheless, response rates to ICI remain below 20-40% and the mechanisms driving ICI response are yet to be understood.

High PD-L1 staining by immunohistochemistry has been the stratification criterion to undergo ICI therapy. However, it presents several pitfalls: first, conflicting evidence has been reported with some responders showing low PD-L1 staining and some non-responders with positive staining (due, for instance, to immune-excluded tumors)^1^. Second, its high heterogeneity, lack of standardization (with different calculation methods (ie. TPS, CPS)), lax positivity threshold, and possible inter-pathologist biased lead to the administration of ICI to almost all candidate patients^2,3^. The tumor mutational burden (TMB, defined as the number of non-synonymous mutations per megabase) has been also considered a candidate biomarker for ICI response, as somatic mutations can act as neoantigens that may favor tumor recognition by infiltrating T-cells during ICI treatment^4,5^. However, TMB was originally calculated through whole-exome sequencing (WES), complicating its clinical implementation due to the need of sufficient input material (not always available in small biopsies) and specific technical equipment and expertise^6^. Despite the attempts to obtain TMB from targeted panels, the discrepancy of the results has prevented its clinical implementation.

But not only the quantity of neoantigens has been associated with ICI response, the quality of the neoantigens (immunogenic) can impact on ICI response. Different sources of high-quality neoantigens have been recently proposed as ICI biomarkers, such as those generated by transposable elements (TE) that can be transcriptionally activated during cancer, which may explain why tumors with low TMB show good immunotherapy responses^7,8^. Among TE, the Pan-Cancer initiative has identified LINE-1 retrotransposons (L1) as being particularly active in certain tumor types^9,10^. Interestingly, most of the tumor-specific TE-chimeric transcripts derive from the LINE class^11^ and analyzing how active L1 can impact ICI response remains a pending question. Parallel to neoantigen generation, antigen processing and presentation is a crucial aspect of antitumor immunity under checkpoint blockade and the repertoire of the Human Leukocyte Antigen (HLA) has recently emerged as a major focus of biomarker discovery efforts^12,13^.

Thus, we have performed a comprehensive characterization of the genomic landscape of advanced lung and bladder primary tumors from patients treated with ICI in first line. First, in lung cancer that remains the leading cause of cancer morbidity and mortality worldwide^14^. Specifically, non-small cell lung cancer (NSCLC) represents 85% of lung cancer cases, and over 40% of patients are diagnosed at advanced stages, when surgical resection is not feasible, leading to a 5-year survival rate of just 9%. Second, in the case of urothelial carcinoma, also referred to as bladder cancer, in which 20-25% of patients present muscle invasive bladder cancer (MIBC) at diagnosis^15^ and, although radical cystectomy is the standard treatment, it only provides a 50% 5-year survival rate^16^. In both cases, ICI have supported a new therapy paradigm representing the gold standard for metastatic NSCLC without an associated targeted therapy^17,18^, and very recently also for metastatic urothelial carcinoma in combination with enfortumab-vedotin^19^, with several clinical trials analyzing its potential in neoadjuvant settings.

Therefore, effective predictive biomarkers represent an urgent need to identify those patients most likely to benefit from ICI therapy. After profiling the genome from NSCLC and MIBC patients treated with ICI in two real-world cohorts, we have identified specific somatic alterations and mutational signatures associated with ICI response. We have additionally evaluated (*) whether non-coding regions impact TMB calculation, (**) the presence of specific HLA-I haplotypes as relevant predictors of response to ICI, and the activation of L1 elements as potential high-quality neoantigens (***).

## RESULTS

### Clinic-pathological characteristics of ICI response

We collected FFPE tumor biopsies at baseline before the start of first line ICI therapy from 34 advanced NSCLC patients with a median age of 63 years old, mostly males and tobacco smokers (Table 1). Both FFPE tumor samples and matching PBMCs from 17 patients underwent WGS (n=34), and RetroTest was applied to 31 FFPE tumor biopsies (Figure 1A). Most patients received single-agent ICI therapy (anti-PD1 pembrolizumab, n=31), while three of them were treated with ICI in combination with chemotherapy at first line. Our cohort consisted mainly of adenocarcinoma histology (n=16). All patients presented high PD-L1 staining (TPS≥50%, median TPS of 80%). Responder patients (R) showed higher PFS (log-rank *p*-value=0.0029, Figure 1B) and OS (log-rank *p*-value=0.011, Supplemental Figure 1A) than non-responder patients (NR). We evaluated whether any clinical variables were associated with improved PFS. Although non-significant, low PD-L1 staining (TPS≤80%) and smoking at the time of diagnosis showed an unfavorable tendency regarding PFS. Neither sex, nor age, reported a statistically significant association with PFS (Figure 1C). Only ICI response showed a statistically significant impact on PFS.

**Figure 1.**
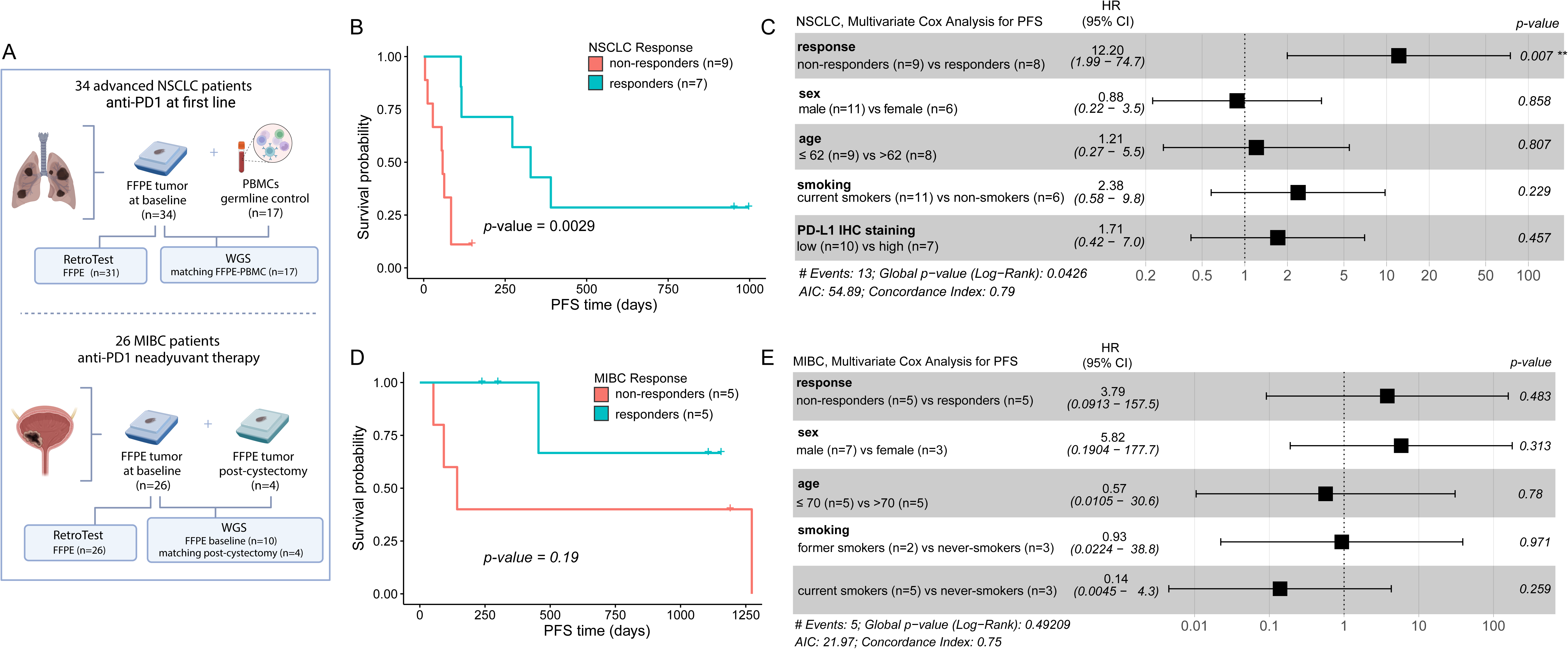
Overview of the NSCLC and MIBC cohorts. **A.** Schematic representation of the two cohorts. For NSCLC patients treated with anti-PD1 at first line, treatment-naive FFPE tumor biopsies were collected for 34 patients and matching PBMCs as germline controls were collected from 17 patients. For MIBC patients who received anti-PD1 neoadyuvant therapy, treatment-naive FFPE tumor biopsies were obtained for 26 patients; plus, matching FFPE tumor samples were obtained post-cystectomy from 4 patients. **B.** Kaplan-Meier curves for PFS with respect to ICI response for NSCLC patients subjected to WGS. Patients were grouped into responders and non-responders to ICI as evaluated by CT scan 3 months after the first anti-PD1 dose. Log-rank test was used to calculate the *p*-value. **C.** Forest plot for the HR of response, sex, age, smoking status and PD-L1 TPS with respect to PFS in NSCLC patients subjected to WGS obtained by multivariate Cox regression. The error bars indicate 95% CI for the HR. **D.** Kaplan-Meier curves for PFS with respect to ICI response for MIBC subjected to WGS. Patients were grouped into responders and non-responders to anti-PD1 evaluated at the time of cystectomy. Log-rank test was used to calculate the p-value. **E.** Forest plot for the HR of response, sex, age and smoking status with respect to PFS in MIBC patients subjected to WGS obtained by multivariate Cox regression. The error bars indicate 95% CI (confidence interval) for the HR.

**Table 1.** Clinicopathological characteristics of the NSCLC cohort.

|  | All patients<br>(n=34) | 3 months after ICI |  | 6 months after ICI |  |  |
| --- | --- | --- | --- | --- | --- | --- |
|  |  | R<br>(n=22) | NR<br>(n=12) | R<br>(n=10) | NR<br>(n=14) | NA<br>(n=10) |
| **Sex** |  |  |  |  |  |  |
| Female | 9 | 4 | 5 | 3 | 6 | 0 |
| Male | 16 | 10 | 6 | 7 | 8 | 1 |
| NA | 9 | 8 | 1 | 0 | 0 | 9 |
| **Age (median)**<br>**(range)** | 63<br>(43 – 78) | 63<br>(51-77) | 60<br>(43-78) | 66<br>(51-76) | 62<br>(43-78) | 59 |
| **Smoking status** |  |  |  |  |  |  |
| **Smoker** | 19 | 11 | 8 | 7 | 11 | 1 |
| Former smoker | 5 | 3 | 2 | 3 | 2 | 0 |
| Never smoker | 1 | 0 | 1 | 0 | 1 | 0 |
| NA | 9 | 8 | 1 | 0 | 0 | 9 |
| **Histological subtype** |  |  |  |  |  |  |
| ADC | 16 | 8 | 8 | 6 | 10 | 0 |
| SCC | 8 | 5 | 3 | 3 | 4 | 1 |
| NA | 10 | 9 | 1 | 1 | 0 | 9 |
| **Tumor stage** |  |  |  |  |  |  |
| IIIC | 2 | 1 | 1 | 1 | 1 | 0 |
| IVA | 6 | 6 | 0 | 4 | 2 | 0 |
| IVB | 9 | 1 | 8 | 1 | 8 | 0 |
| NA | 17 | 14 | 3 | 4 | 3 | 10 |
| **Treatment scheme** |  |  |  |  |  |  |
| ICI | 31 | 20 | 12 | 8 | 14 | 9 |
| ICI-CT | 3 | 2 | 0 | 2 | 0 | 1 |

ADC: adenocarcinoma, CT: chemotherapy, ICI: Immune checkpoint inhibitors, NR: non-responder, R: responder, SCC: squamous cell carcinoma.

In the case of MIBC patients, we collected FFPE tumor biopsies from 26 patients that received ICI as neoadjuvant therapy before radical cystectomy in the context of clinical trials (Table 2). All FFPE baseline treatment-naïve tumor tissues were subjected to RetroTest (n=26), and 10 pre-treatment tumor samples and 4 matching post-cystectomy bladder tissue underwent WGS (Figure 1A). Patients showed a median age of 70 years old. Most were male and tobacco smokers, and most tumors showed urothelial histology (n=9). Response to ICI therapy was evaluated after cystectomy and did not have impact in neither PFS nor OS (log-rank *p*-values 0.19 and 0.99, respectively, Figure 1D, Supplemental Figure 1B). Neither response, sex, age, nor smoking habits reported a statistically significant association with PFS (Figure 1E), confirming the same results as in the original ABACUS clinical^20^.

**Table 2.**
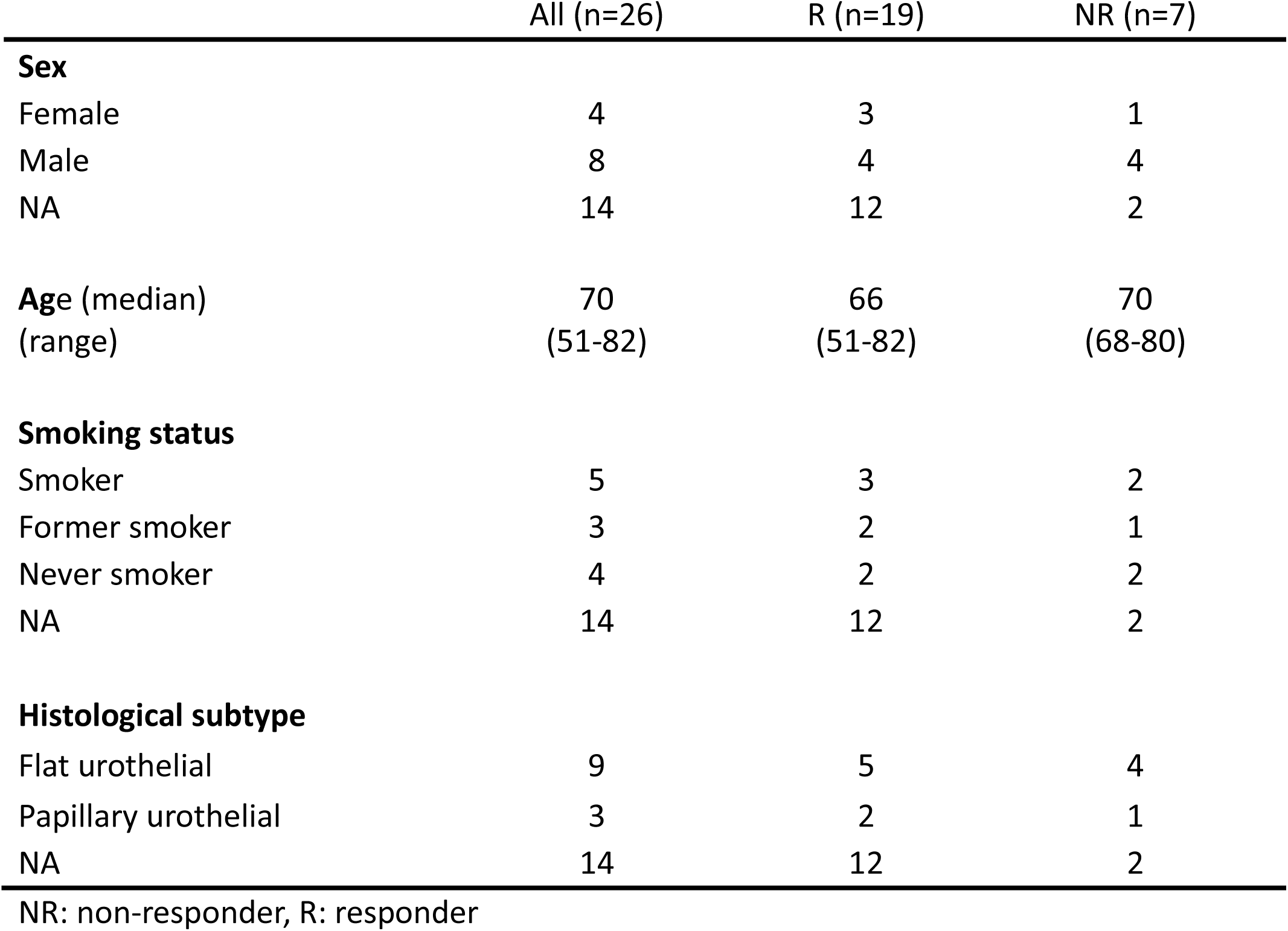
Clinicopathological characteristics of the MIBC cohort.

### NSCLC mutation profiling and TMB in ICI response

We identified an average of 37,432 (9,315 – 104,704) somatic variants (SNVs and INDELS) per sample in a total of 36,063 genes. To identify a list of potential candidate biomarkers for ICI response, we selected genes mutated in, at least, two patients of each group (i.e., mutated in two or more responder patients and not mutated in any of the non-responder patients, or vice versa). We found a total of 1,216 genes exclusively mutated in R patients and 1,748 genes exclusively mutated in NR patients (Supplemental Tables 1, 2). We found that *ATP6V1E1*, *RP11-397E7.1* and *RP11-69I8.2* were the most mutated genes exclusively among R (found in 35.3%), while *FAM63B*, *MYH4*, *OR10D3*, *PROZ*, *RMND1*, *RNU6-743P*, *RP11-122G11.1*, and *RP4-715N11.2* were the top among NR (35.3%).

When focusing on potentially pathogenic variants, we found an average of 92.88 variants per patient (16-316) in a total of 1,316 genes. We did not find differences on TMB, tumor purity, or type of mutation between R and NR (Figure 2A). The most mutated genes with pathogenic variants were *TP53* (82% of the patients), *CSMD3* (47%), and *KRAS* (29%), found both in R and NR. We detected 26 genes harboring pathogenic variants exclusively in R, while 34 genes were exclusively mutated in NR (Supplemental Tables 3, 4). *PIEZO* was found to be exclusively mutated and the most mutated gene among R (50%), while *MT-CO3* gene was exclusive and the most mutated in NR (33.3%).

**Figure 2.**
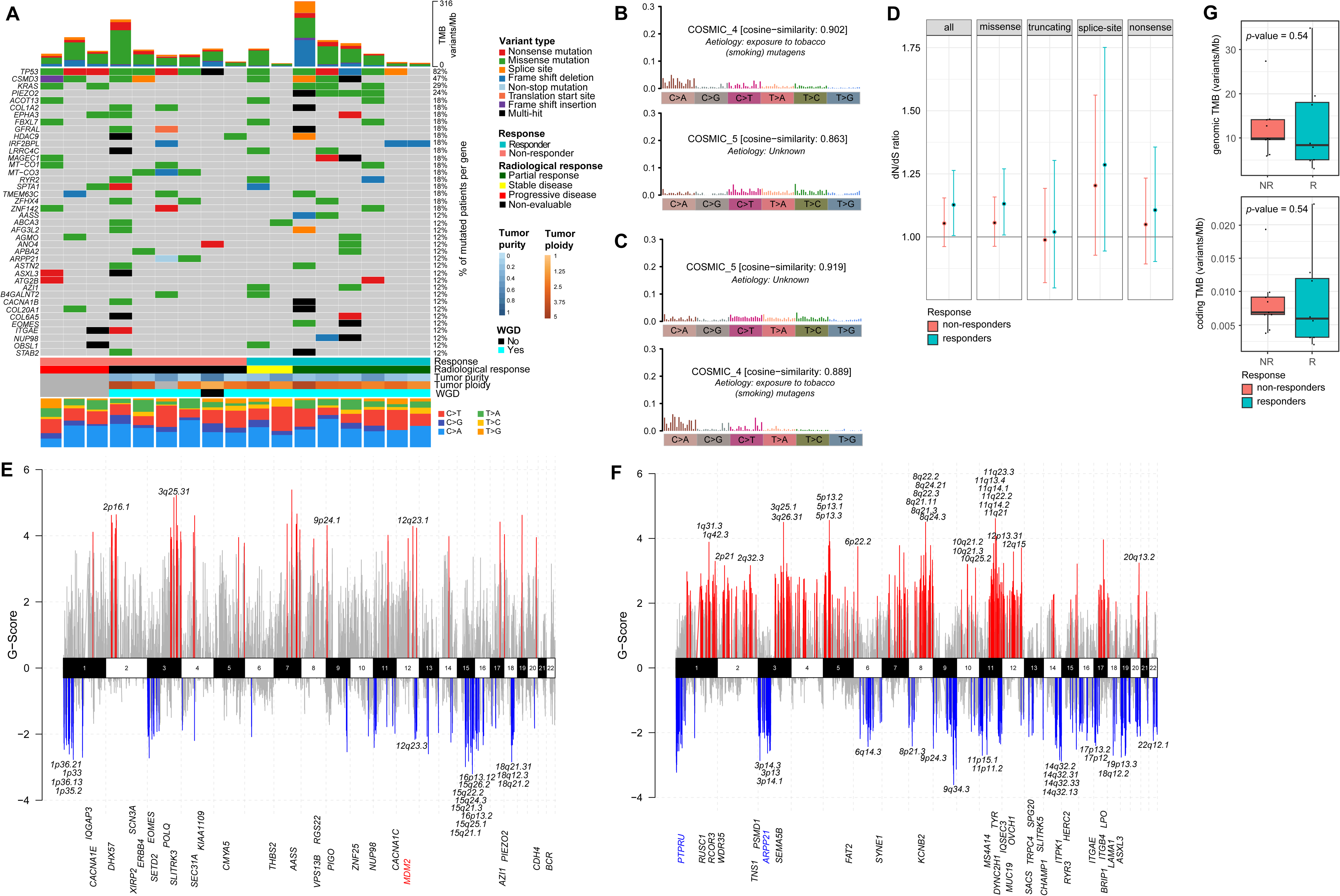
Somatic variants landscape and TMB in NSCLC patients. **A.** Oncoplot showing the top-40 most frequently mutated genes harboring probably pathogenic variants in the cohort of NSCLC patients. **B.** Most common COSMIC mutational signatures for responder patients. **C.** Most common COSMIC mutational signatures for non-responder patients. **D.** Maximum Likelihood Estimation (MLE) of the dN/dS ratio for responder and non-responder patients considering different sets of variants. **E.** Copy number variants profile showing the 66 significantly amplified (red) or deleted (blue) cytobands in responder patients. Those 21 cytobands exclusively altered in responder patients are labelled with their cytoband ID. Genes with probably pathogenic variants found exclusively in responder patients are shown sorted by genomic location. **F.** Copy number variants profile showing the 227 significantly amplified (red) or deleted (blue) cytobands in non-responder patients. From the 179 cytobands exclusively altered in responder patients, the top-50 are labelled with their cytoband ID. Genes with probably pathogenic variants found exclusively in non-responder patients are shown sorted by genomic location. **G.** Boxplots of the TMB calculated considering all variants (left) and of the TMB calculated considering only variants in coding regions (right) with respect to immunotherapy response. Differential p-value was derived by the Wilcoxon rank sum test. The central mark represents the median, with 25th and 95th percentiles at the box, 5th and 95th percentiles at the whiskers and minima and maxima noted by dots.

Afterwards, the COSMIC mutational signatures were extracted from the whole set of variants in both R and NR patients. Signatures COSMIC_4 (“exposure to tobacco (smoking) mutagens”) and COSMIC_5 were the most-common mutational signatures assigned to both R and NR (Figure 2B-C). Based on the dN/dS ratio, we evaluated the mutation selection patterns between responder and non-responder patients. We did not find evidence of positive or negative selection related to response in the different mutation types, although R showed a slight enrichment in missense mutations than NR (Figure 2D).

We next explored the relationship between CNAs and ICI response in our cohort. We detected 13 patients with whole genome duplication (WGD), with an average ploidy of 3.37 (Figure 2A). Moreover, responder patients showed 66 regions with focal CNAs that were statistically significant (FDR *q*-value<5%), of which 21 were exclusively found in R (Figure 2E, Supplemental Table 5). Interestingly, we found two events in *MDM2* with pathogenic variants exclusively found in R. In NR patients, we observed a total of 227 regions with statistically significant CNAs (FDR *q*-value<5%), of which 179 were exclusively present in NR patients (Figure 2F, Supplemental Table 6). The pathogenic variants found exclusively in NR correspond to events overlapping *PTPRU* and *ARPP21*.

Finally, we compared the extent to which TMB can predict ICI response in our real-world cohort following two different approaches: first, we determined the TMB for each tumor sample as “genomic TMB” considering the number total of non-synonymous variants divided by the total genome size. Second, “the coding TMB” was calculated based on the number of non-synonymous variants located exclusively in exonic regions. In both cases, no statistically significant differences were observed between the TMB of NR and R patients (Figure 2G).

### MIBC mutation profiling and TMB in ICI response

We identified an average of 199,221 (117,661-456,401) somatic variants (SNVs and INDELS) per sample in MIBC FFPE tumor tissue at baseline, in a total of 63,167 genes. As before, we selected genes exclusively mutated in at least two patients from each group (R vs. NR) to identify genes potentially associated with ICI response. A total of 5,067 genes were exclusively mutated in R patients, of which 77 genes were mutated in all R (Supplemental Table 7). On the other hand, NR patients showed 721 genes exclusively mutated (Supplemental Table 8). When considering just pathogenic variants, we found an average of 675.5 (137-2,415) variants per sample in baseline samples, in a total of 4,215 genes. The most mutated genes were *TP53* (70%) and *ZFHX4* (60%) (Figure 3A). To identify potential biomarkers for neoadjuvant ICI response, we selected the exclusively mutated genes with pathogenic variants. A total of 614 genes were exclusively mutated in R patients (Supplemental Table 9) and 24 genes were only mutated in NR (Supplemental Table 10). We found that *PKHD1* gene harbored pathogenic variants in all the R patients and that *HEATR5B* and *WDR36* were exclusively mutated in NR patients.

**Figure 3.**
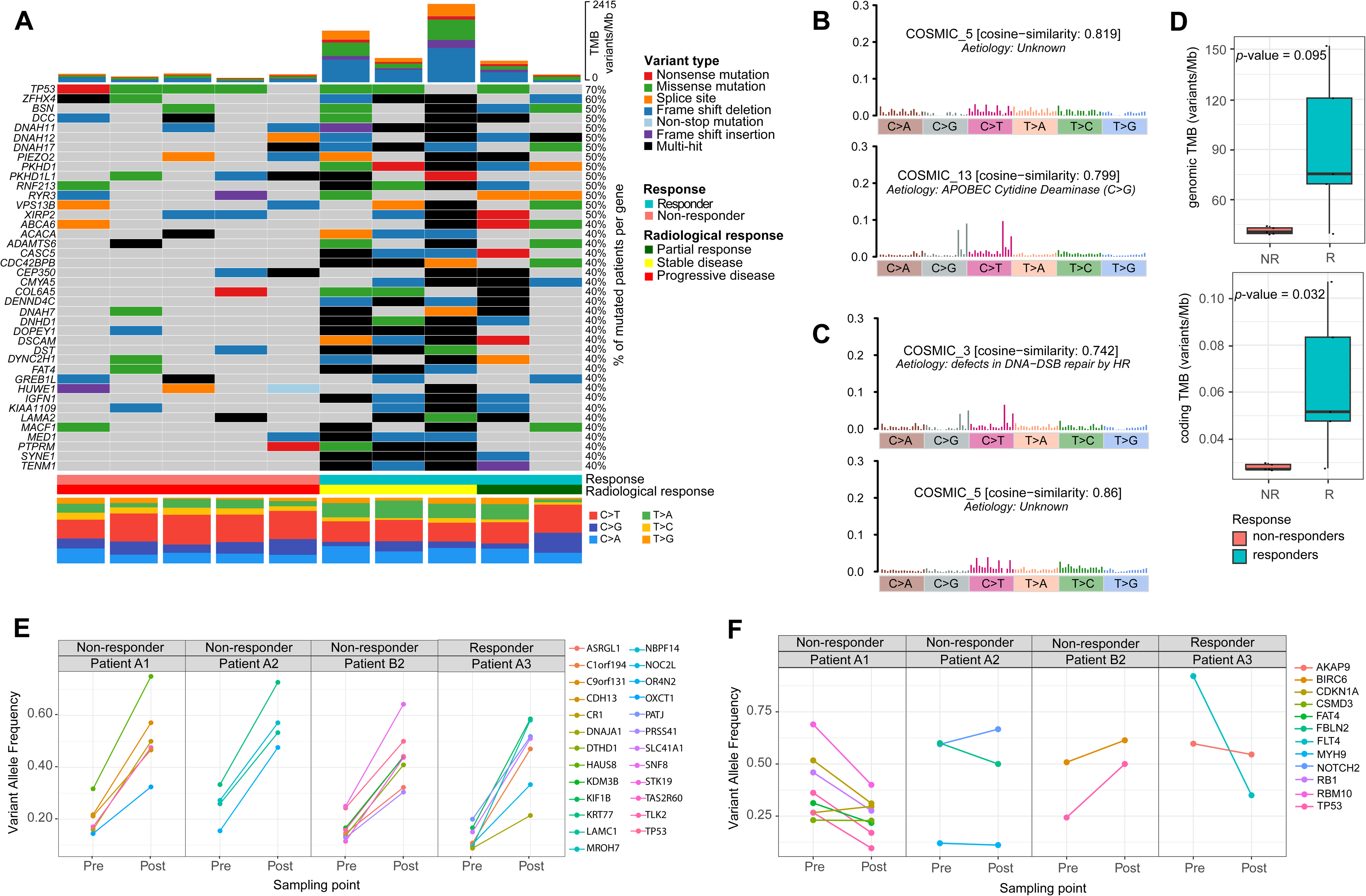
Somatic variants landscape and TMB in MIBC patients treated with neoadyuvant immunotherapy. **A.** Oncoplot showing the top-40 most frequently mutated genes harboring probably pathogenic variants in the cohort of MIBC patients. **B.** Most common COSMIC mutational signatures for responder patients. **C.** Most common COSMIC mutational signatures for non-responder patients. **D.** Boxplots of the TMB calculated considering all variants (“genomic TMB”) and of the TMB calculated considering only variants in coding regions (“coding TMB”) with respect to immunotherapy response. Differential p-value was derived by the Wilcoxon rank sum test. The central mark represents the median, with 25th and 95th percentiles at the box, 5th and 95th percentiles at the whiskers, and minima and maxima noted by dots. **E.** Variant allele frequencies of genes with exonic variants that were shared between a pair of pre-and post-anti-PD1 neoadjuvant therapy. **F.** Variant allele frequency shift of exonic variants present in bladder cancer driver genes.

Using all identified variants, we next extracted the COSMIC mutational signatures in R and NR patients. COSMIC_5 was a common signature to both groups (Figure 3B-C). In contrast, COSMIC_13 (*APOBEC Cytidine Deaminase*) was only found in R (figure 3B), while COSMIC_3 (*defects in DNA−DSB repair by HR*) was exclusively identified in NR (Figure 3C).

We next assessed the predictive potential of TMB calculated following the two different approaches explained above. Interestingly, R patients presented substantially higher TMB compared to NR. However, this difference was only statistically significant when considering coding variants (*p*-value=0.032) (Figure 3D).

### Changes in the mutational profiling during ICI treatment in MIBC

In four MIBC patients, besides the baseline treatment-naïve samples we also obtained a bladder tissue sample from the radical cystectomy after the treatment. Three of the patients were non-responders, showing progressive disease at the time of cystectomy while one showed partial response. We investigated the changes in the mutation profile in the bladder before and after the treatment and reported that post-ICI samples generally have higher SNV count than pre-samples (Supplemental Figure 2A). Next, we focused on the gene identities already altered in the pre-ICI sample. Specifically, we analyzed variant allele frequency (VAF) shifts of the mutations shared between the pairs of matching pre-and post-samples. When considering all variants, we did not find a clear VAF shift pattern between pre-and post-ICI treatment (Supplemental Figure 2B), even when only considering exonic/coding regions (Supplemental Figure 2C). Nevertheless, we detected 25 genes with variants whose VAF substantially increased in post-treatment samples (Figure 3E). When focusing exclusively on MIBC driver genes, we did not find a common pattern, and all the exonic variants that changed their VAF before and after the treatment were patient exclusive. The only gene with VAF changes in two NR patients was *TP53*: one NR patient presented two different *TP53* variants with lower VAF in the post-treatment sample, while another NR harbored a *TP53* mutation positively selected (VAF from 0.25 to 0.5) after ICI therapy (Figure 3F).

### Gene Pathways implicated in ICI response in NSCLC and MIBC

We next performed an enrichment analysis focusing on genes exclusively mutated in only one of the groups (R vs. NR). In the NSCLC cohort, we found that immune related pathways including inflammatory response, and some related to Ras, T-cell receptor/Ras pathway, activation of Ras in B-cells were enriched in R (Supplemental Figure 3A, Supplemental File 1), while the Ras-independent pathway in NK cell-mediated cytotoxicity, neural crest differentiation, PI3K/Akt signaling pathway and FGFR3 and FGFR4 ligand binding and activation pathways appeared enriched in NR (Supplemental Figure 3B, Supplemental File 2). When focusing on pathogenic variants, we observed that R were enriched in several *ERBB4* related pathways such as PI3K events in ERBB2 signaling, signaling by ERBB4 or down regulation of ERBB4 signaling (Figure 4A, Supplemental File 3). In the case of NR, we detected processes related to cell signaling with an important role of integrins, or ECM-receptor interaction (Figure 4B, Supplemental File 4). In addition, adaptive immune system and antigen presentation (folding, assembly, and peptide loading of MHC class-I proteins pathway) were also altered among NR patients. It is worth mentioning that we detected variants affecting other antigen presentation genes more frequently in NR, specifically in genes such as *HERC2*, *PSMD1*, *PSMA6*, and *B2M*. Intriguingly, we realized that SNVs in *B2M* were also amplified in two of the patients.

**Figure 4.**
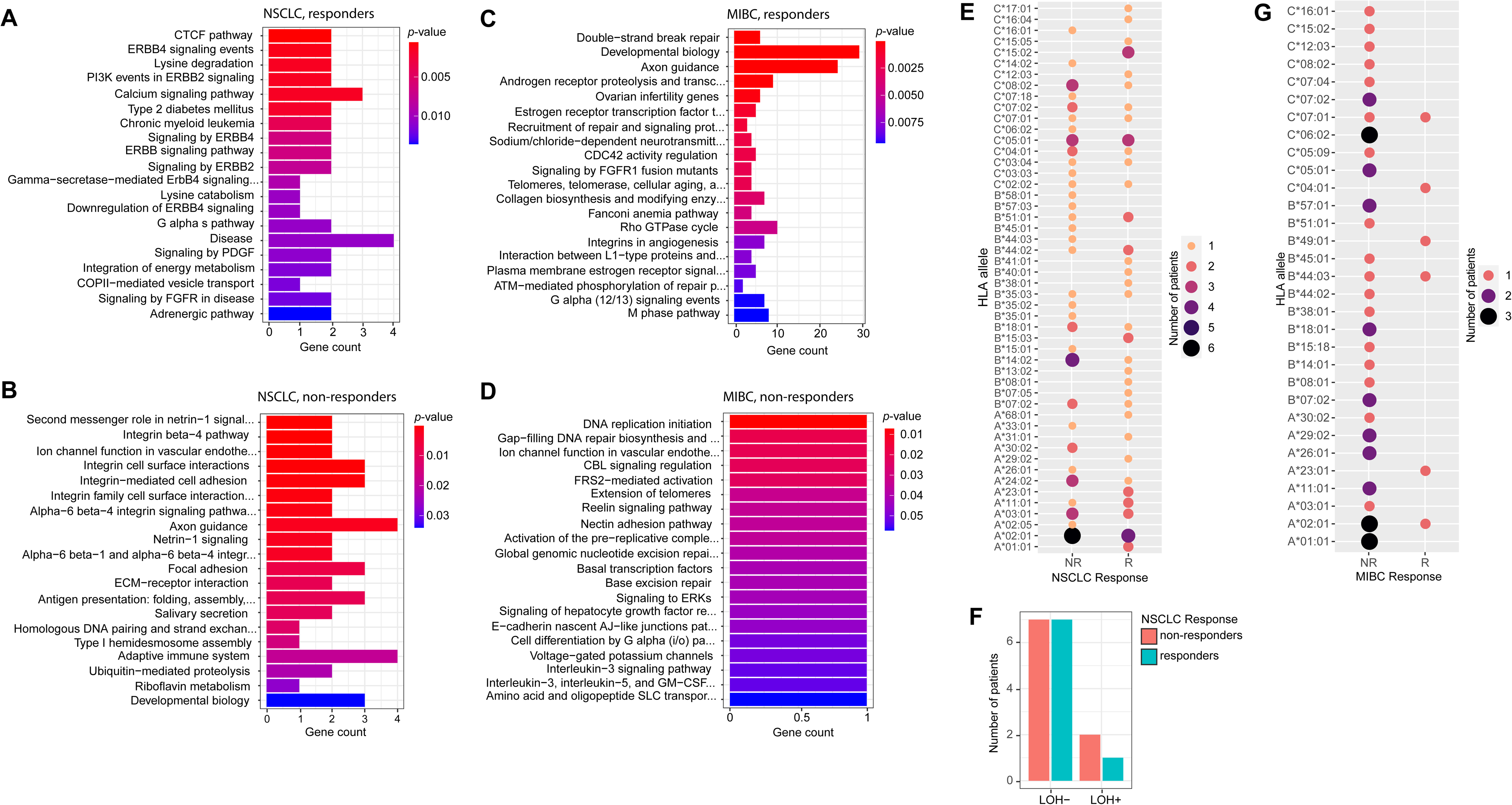
Functional enrichment of somatic variants and HLA typing in responder and non-responder patients. **A.** Barplot of the pathway enrichment analysis based on the 26 genes that contain potentially pathogenic variants exclusive of NSCLC responder patients. **B.** Barplot of the pathway enrichment analysis based on the 34 genes that contain potentially pathogenic variants exclusive of NSCLC non-responder patients. **C.** Barplot of the pathway enrichment analysis based on the 614 genes that contain potentially pathogenic variants exclusive of MIBC responder patients. **D.** Barplot of the pathway enrichment analysis based on the 24 genes that contain potentially pathogenic variants exclusive of MIBC non-responder patients. Enrichment *p*-values were calculated with the Fisher exact test. **E.** HLA-I haplotypes repertoire for NSCLC patients. The bubble size and color indicate the number of patients presenting each specific haplotype. **F.** HLA loss of heterozygosity (LOH) status among responder and non-responder NSCLC patients. A patient was considered positive for HLA^LOH^ if the copy number status estimation for the HLA locus was lower than 0.5. **G.** HLA-I haplotypes repertoire for MIBC patients. The bubble size and color indicate the number of patients presenting each specific haplotype.

In the MIBC cohort, we found no significantly enriched pathways (Supplemental Files 5 and 6), suggesting a high variability in the cohort. When focusing only on pathogenic variants, we found Double-strand break repair pathway, integrins in angiogenesis and interaction between L1-type proteins and ankyrins enriched in R patients (Figure 4C, supplemental file 7). With respect to NR patients, we did not find any statistically significant altered pathways, since all the pathways were only enriched by one gene (Figure 4D, Supplemental File 8), indicating again a huge variability among patients.

### HLA-I Haplotypes repertoire in immunotherapy response

Since cancer biology and ICI response are intimately related to immune processes such as antigen presentation, we next explored the HLA-I locus complexity in both cohorts.

In the NSCLC cohort, the most common haplotype was the HLA-A*02:01, present in four R and in six NR patients (Figure 4E). With respect to ICI response, we found several HLA-I haplotypes that were exclusive to R patients, with HLA-A*01:01, HLA-A*23:01, HLA-B*15:03, and HLA-C*15:02 present in more than one patient. In the case of NR patients, HLA-A*30:02 resulted the only exclusive haplotype, present in two patients, while the alleles HLA-A*02:01 and HLA-B*14:02 were the most common haplotypes among NR, although non-exclusive (Figure 4E). Based on our previous results, we analyzed if HLA loss of heterozygosity occurred more frequently in NR patients, finding that HLA^LOH^ was not statistically significantly higher in NR patients (*p*-value=1, Figure 4F).

For the MIBC cohort, we identified a higher heterogeneity of HLA-I haplotypes in NR patients, without any common haplotype (Figure 4G). Interestingly, when focusing on exclusive haplotypes, HLA-C*06:02 and HLA-A*01:01 appeared exclusively in three NR patients, while HLA-B*49:01 and HLA-A*23:01 were exclusive for R patients (Figure 4G).

### LINE-1 (L1) role on ICI response

Next, we evaluated if L1 activity showed any association with TMB and/or ICI response according to the WGS data in both cohorts. In NSCLC, we did not find a statistically significant correlation between TMB and L1 activation (*p*-value=0.37, Supplemental Figure 4A-B). We also detected that L1 activation was not statistically significantly different between R and NR patients (*p*-value=0.72, Figure 5A). In the case of MIBC, we found a significantly positive correlation between the TMB and L1 activity (*p*-value=0.013, Supplemental Figure 4C-D). Interestingly, L1 activation was clearly higher in R patients, albeit not statistically significant (*p*-value=0.09, Figure 5B).

**Figure 5.**
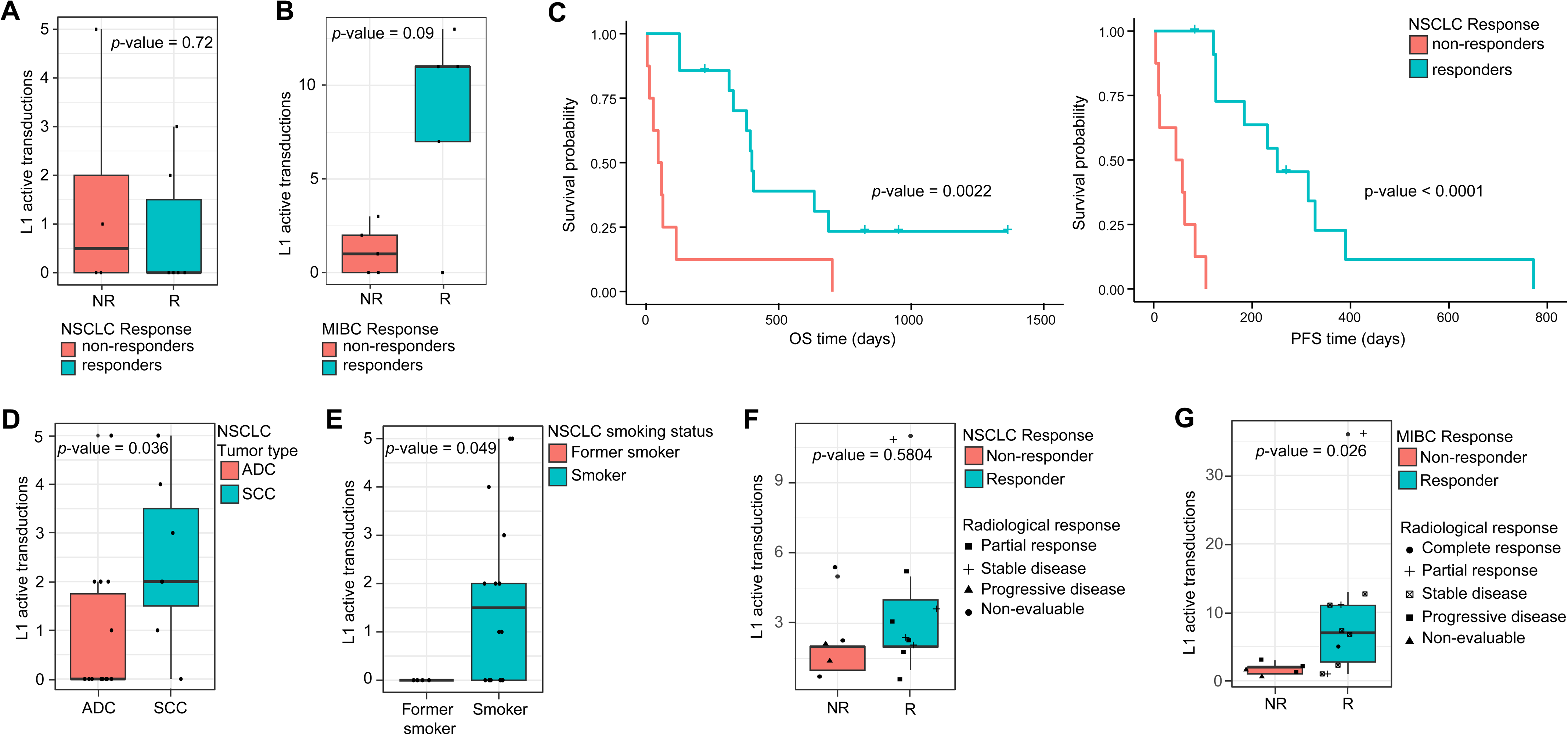
Evaluation of L1 activation measured by RetroTest (L1 active transductions were calculated as the number of TD2 per sample) in NSCLC and MIBC patients treated with ICI. **A.** Boxplot of L1 activation with respect to immunotherapy response in the cohort of NSCLC patients with WGS data (n=17). **B.** Boxplot of L1 activity with respect to immunotherapy response in the cohort of MIBC patients with WGS data (n=10). **C.** Kaplan-Meier curves for OS (left) and PFS (right) according to ICI response in NSCLC patients. The log-rank test was used to derive the p-values. **D.** Boxplot of L1 activation based on the number of transductions according to tumor type. **E.** Boxplot of L1 activity with respect to smoking status (former smokers and current smokers). **F.** Boxplot of the number of transductions in lung cancer patients according to ICI response evaluated at 3 months after the first ICI dose. **G.** Boxplot of the number of transductions in MIBC patients according to response evaluated at the time of cystectomy. For all boxplots, differential p-values were derived by the Wilcoxon rank sum test. The central marks represent the median, with 25th and 95th percentiles at the box, 5th and 95th percentiles at the whiskers, and minima and maxima noted by dots. ADC: adenocarcinoma. NR: non-responder, R: responder, SCC: squamous cell carcinoma

Based on these results, we decided to validate them by specifically determining L1 activity in larger cohorts using RetroTest^21^. We evaluated the impact of ICI response in patients’ prognosis for a larger NSCLC cohort comprising 31 patients (Table 1). First, we confirmed that R patients showed better OS and PFS (log rank *p*-values of 0.002 and <0.0001, respectively, Figure 5C). Moreover, patients with durable response (6 months) also showed better OS (*p*-value=0.002) and PFS (*p*-value<0.0001) (Supplemental Figure 4E). For MIBC, we did not have access to the patients’ follow-up after the cystectomy to perform survival analyses. When analyzing L1 retrotransposition in these NSCLC patients, we found that patients with squamous tumors showed statistically higher L1 activity than those with adenocarcinomas (*p*-value=0.032, Figure 5D), while current smokers showed higher activation than former smokers (*p*-value=0.049, Figure 5E). Finally, when we evaluated L1 activity with respect to ICI response, we detected that 45.16% of patients showed L1 activation. In addition, we noticed that most patients with high L1 activation were R, although not reaching statistical significance (*p-*value=0.5804, Figure 5F).

In the case of L1 retrotransposition in a larger MIBC cohort (n=26) (Table 2), we found that 57.7% of the patients presented L1 activation. We observed that all patients with high L1 activation were R (*p*-value=0.026, Figure 5G), thus following the same tendency observed in NSCLC, but this time reaching statistical significance, hence highlighting the power of L1 activation as predictive biomarker for ICI response.

## DISCUSSION

ICI have achieved a milestone in advanced cancer treatment, although the identification of response predictive biomarkers is still limited. In fact, it has become the first-line treatment for advanced NSCLC^18^, recently approved for MIBC in combination with enfortumab-vedotin^19^, and as neoadjuvant treatment in NSCLC^22^. In this scenario, the lack of effective predictive biomarkers of response represents a key challenge in clinical practice. In this study, we provide a wide genomic characterization for NSCLC and MIBC treated with ICI as first-line therapy.

Importantly, none of the analyzed clinical characteristics showed a statistically significant association with patient prognosis, neither in NSCLC, nor in MIBC, highlighting the need of reliable and robust genomic biomarkers for oncological patient management.

Our results confirmed that key cancer driver genes are often mutated in both R and NR patients, such as *TP53*, the most mutated gene for both tumor types, *KRAS* and *MT-CO3* in NSCLC, and *ZFHX4* in MIBC. Nevertheless, we also identified specific mutations, which could potentially serve as predictive biomarkers for ICI response, such as *PIEZO2* and *MT-CO3*, specific of R and NR respectively in NSCLC patients, while *PKHD1* and *HEATR5B/WDR36* appeared exclusively in R and NR respectively in MIBC.

Moreover, by exploring the active mutational signatures in these cohorts, we found that COSMIC_4 (tobacco exposure) and COSMIC_5 were common in all NSCLC patients. The presence of the COSMIC_4, with 94% of ever-smoker patients, is expected and in agreement with what has been described for NSCLC^23,24^. In relation to MIBC, we identified APOBEC cytidine deaminase signatures (COSMIC_13) exclusively among R patients. A correlation between APOBEC-mediated mutagenesis and ICI effectiveness has already been described in MIBC^25^, with several studies linking APOBEC mutagenesis to neoantigen generation^26,27^. Hence, the presence of this feature could not only potentially represent a biomarker for ICI response, but also underlie the mechanisms behind this process in MIBC patients.

In relation to TMB, we detected no statistically significant association between the TMB and ICI response in NSCLC. In contrast, MIBC R patients showed a significantly higher TMB than NR patients when considering the coding regions. These results appear to indicate that WES-based estimates may be sufficient, as our WGS TMB metrics did not contribute to a higher predictive power. In any case, it should be mentioned that TMB was only found to be an effective biomarker of ICI response in MIBC.

With the aim of digging deeper into the mechanisms that could underlie ICI response, we explored the molecular pathways enriched in the sets of exclusively mutated genes in R and NR patients. We found pathways related to inflammatory response and to Ras dependent immune molecular genes exclusively mutated in NSCLC R*. Ras* family has been widely studied as a key regulator of the activity of multiple downstream signaling pathways, including cell proliferation^28^. In particular, oncogenic *RAS* mutations have been described as promoting immune evasion by favoring the expression of PD-L1 in tumor cells^29,30^, which could partly explain why ICI therapy is more effective in these patients. Similar to previous studies, we also found mutations in ERBB related pathways in R. *ERBB4* alterations have been described as hampering immunotherapy response in NSCLC^31^, thus promoting PD-L1 expression^31–33^. In addition, we described its co-mutation with *MDM2* gene in one R patient, whose inhibition was reported to increase MHC class I and II expression in a *TP53* dependent manner, its main inhibitor, and, hence, to promote neoantigen recognition and T-cell infiltration^34,35^. Accordingly, there are different clinical trials targeting *MDM2* in combination with immunotherapy^34,36^. Finally, genes belonging to CTCF pathway, closely related to chromatin remodeling and epigenetics, presented pathogenic variants exclusively in R. Notably, CTCF has been described as an inhibitor of lncRNA responsible of reducing immunotherapy response^37^.

Regarding NR patients, we found an enrichment of FGFR3 ligand binding and activation pathways. Intriguingly, *FGFR3* mutations have been recently described to drive T-cell-depleted microenvironment in bladder cancer, attenuating the response to ICI in metastatic MIBC patients^38^. Moreover, we discovered an important enrichment on neural crest differentiation, which is in line with recent research pointing towards a neuronal regulation of immune response mediated by innate lymphoid cells (ILCs)^39–41^. This is noteworthy considering the emerging field of cancer neuroscience and how ICI response could be strongly modified by neuromodulation strategies^42,43^. We also detected exclusive alterations affecting cell-surface and cell-cell interaction components, such as integrins, and MHC class-I. In this regard, the blockage of Netrin-1, a member of one of the pathways in which these integrins are involved, has been associated to a better response to ICI^44^, while MHC class-I components impairment can hamper neoantigen presentation and have been proposed as tumor resistance processes^45–47^. In addition, we identified variants more frequently in NR affecting several genes related to antigen presentation, such as *HERC2*, *PSMD1*, *PSMA6*, and *B2M* (*B2M* appeared also amplified in two of the patients). Our observations also indicated that an impaired HLA-I complex affects the ICI response. These results are in line with previous studies reporting recurrent inactivation of *B2M* in lung cancer, together with a down-regulation of the HLA-I complex, leading to an abnormal immunosurveillance in lung cancer^48^. Thus, our results confirm a molecular resistance to the ICI therapy by impairing the presentation of immunogenic neoantigens.

In MIBC patients, Double Strand Break Repair system alterations were found in R patients. Concordantly, previous studies have shown that these impairments can cause genomic instability^49,50^, PD-L1 expression and TMB upregulation, and immunotoxicity through cGAS-STING stimulation, which favor interferon expression and lymphocytes infiltration^50–53^. Furthermore, the impairment of this repair system may also trigger neoantigen generation^50,54^, favoring ICI response. In relation to this, *ATM* mutations have also been reported in other NSCLC cohorts in patients who responded to ICI therapy^24^ while its inhibition has been described to enhance cancer immunotherapy by promoting cGAS/STING activation^55^. Among NR patients, we found a huge mutational variability, which likely indicates the activation of a mixture of molecular pathways preventing the response in a neoadjuvant scenario.

We additionally explored features related to the tumor microenvironment by analyzing the HLA-I haplotype repertoire. In each cohort, we found several common HLA haplotypes but also identified several exclusive haplotypes to both R and NR subgroups. Stratifying patients based on this genetic background could potentially guide ICI decision making. Specific HLA-I alleles have been already related to tumor infiltrating lymphocytes cytotoxicity activation after antigen presentation^56^, and the presence of HLA-A*03 alleles has been recently suggested as a biomarker of poor response to ICI^57^. Indeed, this specific allele was found more frequently within our NR patients. HLA-A*02:01 allele was the most frequent in our cohort of MIBC, especially in NR, and has been recently associated with good prognostic in pancreatic cancer^58^. Therefore, these novel results support the need to further study the HLA haplotypes as potential predictive biomarkers for ICI response.

Finally, recent studies have reported that the activation of TEs, such as L1, can enhance ICI response by inducing inflammation and generating immunogenic neoantigens^8,11,59^. However, the relation between L1 activation and immunotherapy response in real-life ICI-treated patients is yet to be evaluated. Based on this and on our first results showing a higher tendency of L1 activation in tumors with higher TMB resulting in R patients, we decided to further validate the possible association between L1 activation and ICI response in extensive cohorts. In general, we found that half of the patients presented L1 activation at baseline, varying slightly among the NSCLC and MIBC patients. In NSCLC, a significant positive association was found between smoking and L1 activity, finding an association with squamous tumors, intimately linked to smoking habits. Interestingly, we detected a clear tendency in both NSCLC and MIBC, showing a higher L1 activation in R compared to NR patients, although only statistically significant in MIBC. To the best of our knowledge, this is the first time that the capacity of L1 activation was demonstrated to predict ICI response at baseline.

In summary, our findings highlight that while several cancer driver genes, such as *TP53*, are commonly mutated in both R and NR patients, but specific mutations were also as potential predictive biomarkers for ICI response. The analysis of active mutational signatures revealed interesting patterns that may relate to treatment efficacy. In this context, the relationship between tumor mutational burden (TMB) and ICI response resulted evident in MIBC patients but not for NSCLC. The examination of molecular pathways also indicated that certain mechanisms, such as immune evasion driven by RAS mutations and PD-L1 expression, play a crucial role in treatment effectiveness. Findings regarding the activation of L1 suggest a potential link between induced inflammation and response to ICI, opening new avenues for developing additional biomarkers. Ultimately, our results underscore the importance of a genomic approach in cancer management and the need for further studies to validate these biomarkers and better understand the underlying mechanisms affecting response to ICI therapy. This will not only improve clinical management for patients but also contribute to a more personalized approach in cancer treatment.

## METHODS AND MATERIALS

### Patients’ cohorts, response and survival

We included a total of 34 advanced NSCLC patients collected from Hospital Álvaro Cunqueiro (Complexo Hospitalario Universitario de Vigo, CHUVI, Spain) and Complexo Hospitalario Universitario de Santiago (CHUS, Santiago de Compostela, Spain). All patients were treated with ICI at first line, either alone or in combination with chemotherapy (Table 1). We characterized the mutation profile of 17 patients using whole genome sequencing (WGS) and evaluated L1 retrotransposition in 31 patients. Treatment-naive formalin-fixed paraffin-embedded (FFPE) primary tumor biopsies were collected and peripheral blood mononuclear cells (PBMCs) from the same NSCLC patients, to be used as germline control. Immunotherapy response was evaluated by computed tomography scan 3 and 6 months after the first dose. Durable response was evaluated according to the response at 6 months.

Furthermore, we included 26 MIBC patients involved in various clinical trials for neoadjuvant ICI treatment evaluation at Hospital “12 de Octubre” (Madrid, Spain) (Table 2). We analyzed a sub-cohort of 10 patients from the ABACUS clinical trial^20^ using WGS and the 26 patients for L1 activation. FFPE primary tumor biopsies were obtained and response to ICI was evaluated at cystectomy. Post-ICI FFPE tumor samples were also acquired at cystectomy for 4 out of the 10 patients subjected to WGS.

In both cohorts, patients with stable disease, partial response, and complete response were classified as responders (R), while non-evaluable patients (*exitus*) or patients with disease progression at the time of evaluation were considered non-responders (NR). Overall survival (OS) was defined as the time from treatment initiation to the time of death, while progression-free survival (PFS) was defined as the time from treatment initiation to the time of progression or death, whichever is earlier. The study was conducted with appropriate authorization from the Galician Regional Research Ethics Committee (2019/046) and from the Ethics Committee from Hospital “12 de Octubre” (17/094) following the Helsinki Declaration of 1975 and all patients signed informed consent approving their participation.

### Sample preparation and WGS

We isolated genomic DNA from FFPE samples from NSCLC and MIBC tumors using QIAamp DNA FFPE Tissue kit (Qiagen). PBMCs from NCLC patients were obtained by Ficoll density gradient from a blood sample drawn before the first ICI dose (baseline). We extracted DNA from PBMCs with the QIAamp DNA Blood Mini kit (Qiagen), according to manufacturer’s protocol, as germline control. We checked DNA quality using Qubit dsDNA BR Assay Kit in Qubit 4.0 (Thermo Fisher) and quantified the total yield and gDNA ScreenTape Analysis in a 4200 TapeStation system (Agilent Technologies) to assess the DNA integrity. Samples were sequenced in an external service (Macrogen) with Illumina Truseq Nano DNA Library in a NovaSeq6000 (Illumina) at 150bp PE.

### WGS data processing

We assessed the quality of raw WGS data with *FastQC*^60^. We next mapped the sequencing reads from both tumor and PBMC samples to the GRCh37/hg19 reference genome using the Burrows-Wheeler Aligner (*BWA-mem* v0.7.17)^61,62^. Afterwards, we used *Samtools* v1.9^63^ to sort the aligned reads and to index the generated bam files. Duplicated reads were marked with Picard *Bammarkduplicates*^64^. Finally, we B*ase Quality Score Recalibration* (BQSR) using the Genome Analysis Tool Kit (GATK) v.4.1.7.0^65^ to correct quality score bias, following the best practices guidelines^66^.

### SNVs and INDELs calling

Before variant calling, we estimated cross sample contamination using GATK’s *GetPileupSummaries* and *CalculateContamination*, as well as the genomAD resource obtained from the GATK bundle (https://console.cloud.google.com/storage/browser/gatk-best-practices/somatic-b37;tab=objects?prefix=). For NSCLC, we performed cross sample contamination in paired-sample mode, using the PBMCs data as germline control, while for pre-treatment MIBC samples it was performed in tumor-only mode.

Next, we used *Mutect2* v0.7.17^67^ for calling SNV and indels. For NSCLC samples, the tumor-normal matched mode was used, with PMBCs data as normal. For pre-treatment MIBC samples, we performed variant calling in tumor-only mode. For the post-treatment samples analysis, we used the matching pre-treatment sample as control. We next applied *FilterMutectCalls* with standard parameters to remove low quality variants, considering the estimates from both *CalculateContamination* and *LearnReadOrientationModel*, which identifies paraffin related artifacts. Those variants selected by *FilterMutectCalls* in at least two patients were considered artifacts and discarded from further analyses. The remaining variants were finally annotated with the Ensembl Variant Effect Predictor (VEP) v100.2^68^. We identified likely pathogenic variants according to SIFT^69^ and Polyphen^70^ scores and used *vcf2maf* v1.6.21^71^ to transform the VCF files to MAF format. We used the *mafTools* R package v2.16.0^72^ for downstream analyses and visualization purposes.

### SNVs and INDELs custom filtering

To further remove low quality variants and potential artifacts for mutational signatures extraction and functional enrichment, we additionally removed mutations occurring in the following genome conflicting regions according to UCSC Genome Browser: (i) sites present in the NCBI track dbVar Curated Common SVs Conflicts with Pathogenic (DbVarConflict), which includes common structural variants that overlap with structural variants with ClinVar annotation^73^; (ii) variants present in the regions of the ENCODE blacklist subtrack, which contains a set of problematic regions due to a high ratio of multi-mapping to unique mapping reads and high variance in mappability due to repetitive elements^74^; (iii) sites present in the database of fragile sites in human chromosomes (HumCFS)^75^; and (iv) variants overlapping with the Genome-In-A-Bottle (GIAB) giabCallConflict track set, that contains a set of regions where it is difficult to make a confident call, due to low coverage, systematic sequencing errors, and local alignment problems^76^.

### TMB and the ratio of non-synonymous vs. synonymous substitutions (dN/dS)

We determined the TMB of each tumor sample as follows: (i) the genomic TMB was calculated as the 10-base logarithm of the number total non-synonymous variants divided by the total genome size, while (ii) the coding TMB was calculated as the 10-base logarithm of the number of non-synonymous variants located in exonic regions divided by 10^^6^.

Regarding dN/dS analysis and MIBC post-therapy variant analysis, we decided to reduce even more the noise from low frequency and unreliable calls, so we removed variants with less than 10 supporting reads in either the tumor or the corresponding PBMC sample, variants that were called in the PBMC sample, and variants with less than 0.075 variant allele frequency (VAF) in the tumor sample. Additionally, as variant calling in MIBC pre-treatment samples was performed in tumor-only mode, we discarded variants with a VAF greater than 0.45 to rely only on subclonal variants.

Afterwards, these filtered variant sets were used to estimate the dN/dS score for each tumor sample to infer selection pressure in protein-coding genes through dNdScv, which applies a maximum likelihood approach to quantify selection patterns^77^.

### Single-Base Substitution (SBS) signature extraction & assignment

To study mutational signatures, we used the *trinucleotideMatrix* function from *maftools* to extract the immediate bases flanking the mutated site and to classify them into 96 classes depending on the combination of the trinucleotides. Then, we applied *extractSignatures* function to decompose the matrix into *n* signatures. The number of signatures was chosen based on the value at which Cophenetic correlation drops significantly (which can be calculated through *estimateSignatures* and *plotCophenetic* functions). Finally, we used *compareSignatures* function cosine similarity to detect the best similarity to COSMIC database known signatures^78^.

### CNV calling

We identified the copy number variants (CNV) of NSCLC tumor samples with the purity and ploidy estimator (*PURPLE*)^79^ in the paired-sample mode under default parameters, using tumor and PBMCs bam files together with the SNV and indel calling as input. We determined different CNV regions between responder and non-responder patients, together with the genes contained in each aberrant region with *GISTIC2.0*^80^, using the CNV segments in each sample. The identification of differentially amplified/deleted regions was based on a G-score assigned to each of them, depending on their amplitude and their alteration frequency across the set of samples of each group. A *q*-value false discovery rate of 25% was set to call a significant region and then the window with the greatest amplitude and frequency was considered the “peak” of the region. We determined the boundary of each peak by the *RegBounder* algorithm implemented in *GISTIC2.0*, with a 95% confidence level to include target genes within each region^80^.

### Enrichment analysis

We performed an enrichment analysis using *enrichR* R package^81,82^, an interface to the Enrichr database hosted at https://maayanlab.cloud/Enrichr/. This tool assesses whether our differentially mutated gene sets were enriched in specific biological/molecular functions by measuring the statistical significance (Fisher exact test) of the overrepresentation of a gene set in a specific list of genes. We selected the following databases: Human MSigDB collections (including MSigDB_Hallmark_2020, MSigDB_Oncogenic_Signatures, and MSigDB_Computational)^83,84^, BioPlanet_2019^85^, KEGG_2019_Human^86,87^, WikiPathways_2019_Human^88,89^, GO_Molecular_Function_2018, and GO_Biological_Process_2018^90,91^, ChEA_2016^81^ and TRANSFAC_and_JASPAR_PWMs^92^.

### HLA-I Haplotyping and HLA LOH analysis

To identify HLA-I haplotypes we applied *LILAC*^93^, a tool which determines HLA-I germline alleles; by aligning the sequencing reads to HLA-A, HLA-B and HLA-C regions which are present in the IMGT/HLA database^94^. Following their recommended pipeline, we first performed an “elimination” phase, removing allele candidates that are clearly not present in the sample, and an “evidence” step, which selects the best solution based on the observed reads amongst the whole set of possible HLA-I alleles. For the NSCLC cohort, we used both tumor and PMBCs bam files as input, together with the somatic variants (SNVs, INDELS, and CNA). For the MIBC cohort, we used tumor bam files and somatic variants (SNVs and INDELS). We also determined Loss of Heterozygosity (LOH) events affecting HLA haplotypes (HLA^LOH^) for NSCLC patients when the copy number estimated by *PURPLE*^79^ (see Methods section 2.6) was less than 0.5.

### LINE-1 retrotransposition rate quantification by RetroTest

To quantify the amount of LINE-1 (L1) somatic insertions, we applied the RetroTest targeted sequencing method^21^, based on the use of probes that capture the unique sequence downstream of the 124 most active L1 elements of the human genome^9^. RetroTest library construction and bioinformatic processing were performed following the recommended approach^21^.

### Statistical analyses

We performed survival analyses using OS and PFS, with censoring at the date of the last follow-up if a patient was alive and without disease progression. We combined clinical variables (sex, age, smoking habits and PD-L1 staining) in a multivariate Cox proportional hazards ratio (HR) model with PFS as response variable, with the *coxph* function from the *survival* v3.5.7 R package. We estimated survival curves using the Kaplan-Meier method and compared them with the log-rank test. We assessed the significance between the TMB, the L1 insertion rate and the presence of HLA^LOH^ between responder and non-responder patients with the non-parametric Wilcoxon rank sum test (*p*-value threshold of 0.05). For multi-testing correction, we computed FDR values from nominal *p-values* using the Benjamini-Hochberg method.

## Acknowledgements

Authors thank all the enrolled patients and their families. Samples were collected and stored by the Galicia Sur Health Research Institute (IIS Galicia Sur) Biobank (registry B.0000802). This research project was made possible through the access granted by the Galician Supercomputing Center (CESGA) to its supercomputing infrastructure. The supercomputer FinisTerrae III and its permanent data storage system have been funded by the Spanish Ministry of Science and Innovation, the Galician Government and the European Regional Development Fund (ERDF).

This work was supported by the Instituto de Salud Carlos III (ISCIII) and the European Social Fund (“Investing in your future”) (PI19/01113) and the Spanish Association Against Cancer Scientific Foundation (IDEAS19122MART). J.B.I. was supported by the Spanish Association Against Cancer Scientific Foundation (PRDCR19007BREA_001). A.O. was supported by predoctoral fellowship from the Galician Innovation Agency, Xunta de Galicia (ED481A-2020/214). M.M.F was previously supported by the Spanish Association Against Cancer Scientific Foundation (INVES207MART) and is currently supported by the Miguel Servet program (CP20/00188) from the Instituto de Salud Carlos III (ISCIII) and the European Social Fund (“Investing in your future”). M.G-G. is currently supported by a postdoctoral fellowship from the Galician Innovation Agency, Xunta de Galicia (IN606B-2024/014).

## DATA AVAILABILITY

Whole Genome Sequencing data from the NSCLC and MIBC cohorts are deposited into the Sequence Read Archive (SRA) repository under the following BioProject IDs: PRJNA1146979 and PRJNA1144206, respectively. RetroTest pipeline is publicly available at: https://gitlab.com/mobilegenomesgroup/RETROTEST.

## AUTHOR CONTRIBUTIONS

JB-I, MG-G, AO, MM-F contributed to the design of the study. MEL-Q, LL, LJM, CG-B, IA, LM, JMP contributed to the acquisition of the data set. JB-I, MG-G, AO, JMA contributed to the analysis of data and their interpretation. JB-I and MM-F drafted the paper. JB-I, MG-G, AO, JMA, JMP, MM-F contributed to their interpretation and manuscript editing. All authors reviewed and revised the work and approved the final draft for submission.

---

**Supplemental figure 1. A.** Kaplan-Meier curves for OS with respect to ICI response for NSCLC. Patients were grouped into responders and non-responders to ICI as evaluated by CT scan 3 months after the first anti-PD1 dose. Log-rank test was used to calculate the p-value. **B.** Kaplan-Meier curves for OS with respect to ICI response for MIBC. Patients were grouped into responders and non-responders to ICI as evaluated at cystectomy. Log-rank test was used to calculate the p-value.

**Supplemental figure 2. A.** Mutational burden estimates of pre-and post-neoadyuvant immunotherapy in MIBC samples, showing the total counts per sample of INDEL and SNV variants. **B.** Scatterplot comparing the VAF of all the variants that are shared between the pre-and post-neoadyuvant immunotherapy in MIBC patients. **C.** VAF shift between exonic variants that are shared between the pre-and post-neoadyuvant immunotherapy in MIBC patients. VAF shift was calculated by subtracting the VAF of the pre-ICI sample to the VAF of the post-ICI sample. Variants whose VAF increases after neoadyuvant ICI are shown in the red scale, while variants with lower VAF after neoadyuvant ICI are shown in the blue scale.

**Supplemental figure 3. A.** Barplot of the pathway enrichment analysis based on the 1,216 genes that are exclusively mutated in NSCLC responder patients. **B.** Barplot of the pathway enrichment analysis based on the 1,748 genes that are exclusively mutated in NSCLC non-responder patients. **C.** Barplot of the pathway enrichment analysis based on the 5,067 genes that are exclusively mutated in MIBC responder patients. D. Barplot of the pathway enrichment analysis based on the 751 genes that are exclusively mutated in MIBC non-responder patients.

**Supplemental figure 4. A.** Scatterplot for the correlation between the TMB calculated with all genomic variants and the LINE-1 activation measured by RetroTest for NSCLC patients. **B.** Scatterplot for the correlation between the TMB calculated with exonic variants and the LINE-1 activation measured by RetroTest for NSCLC patients. **C.** Scatterplot for the correlation between the TMB calculated with all genomic variants and the LINE-1 activation measured by RetroTest for MIBC patients. **D.** Scatterplot for the correlation between the TMB calculated with exonic variants and the LINE-1 activation measured by RetroTest for MIBC patients. **E.** Kaplan-Meier curves for OS (left) and PFS (right) according to durable response to ICI in NSCLC patients. The log-rank test was used to derive the p-values.

