## Supplementary figures and images for "Genomic Insights Guiding Personalized First-Line Immunotherapy Response in Lung and Bladder Tumors"

### Supplemental Figure 1

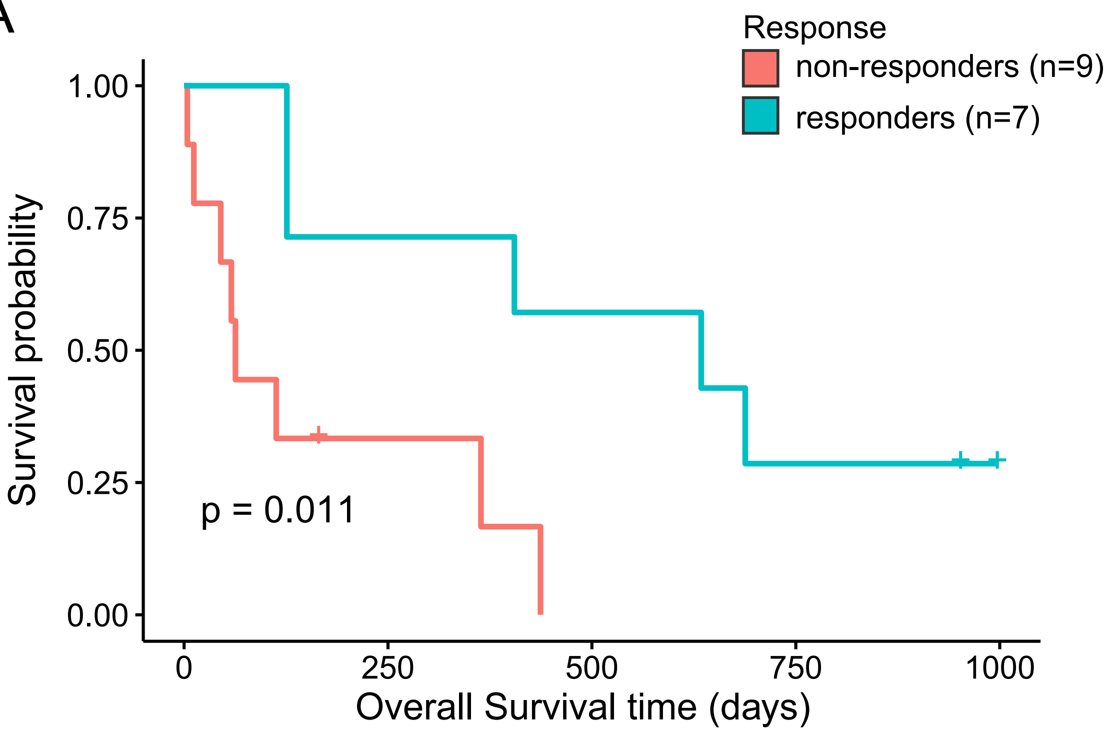

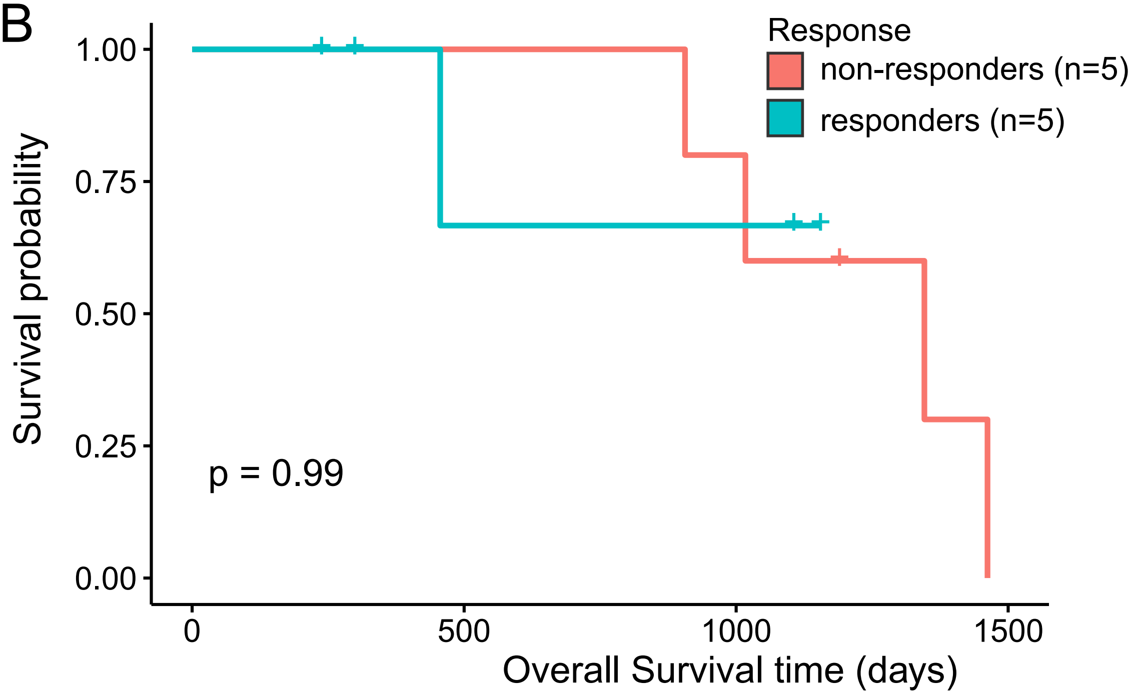

### Supplemental Figure 2

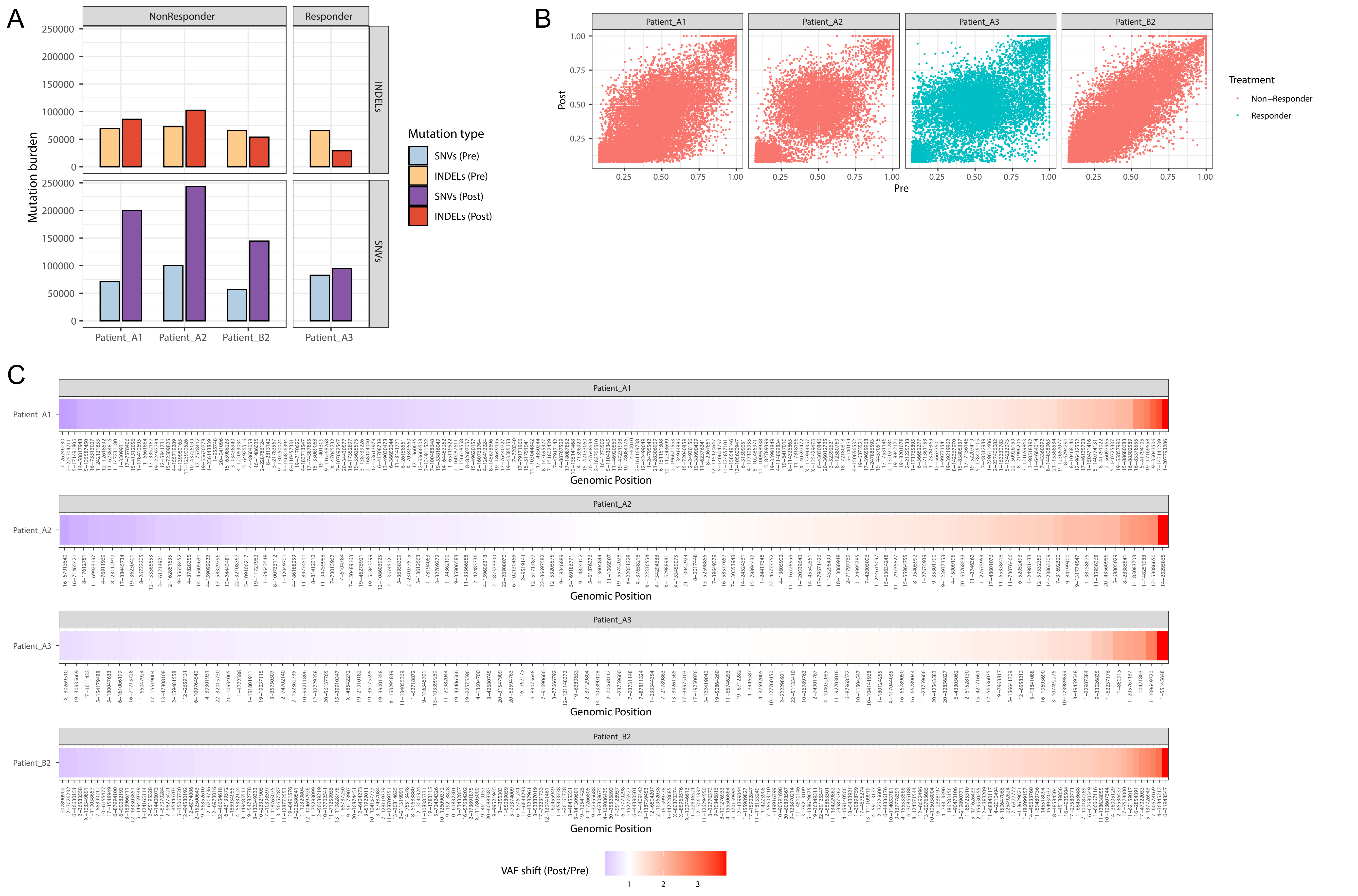

### Supplemental Figure 3

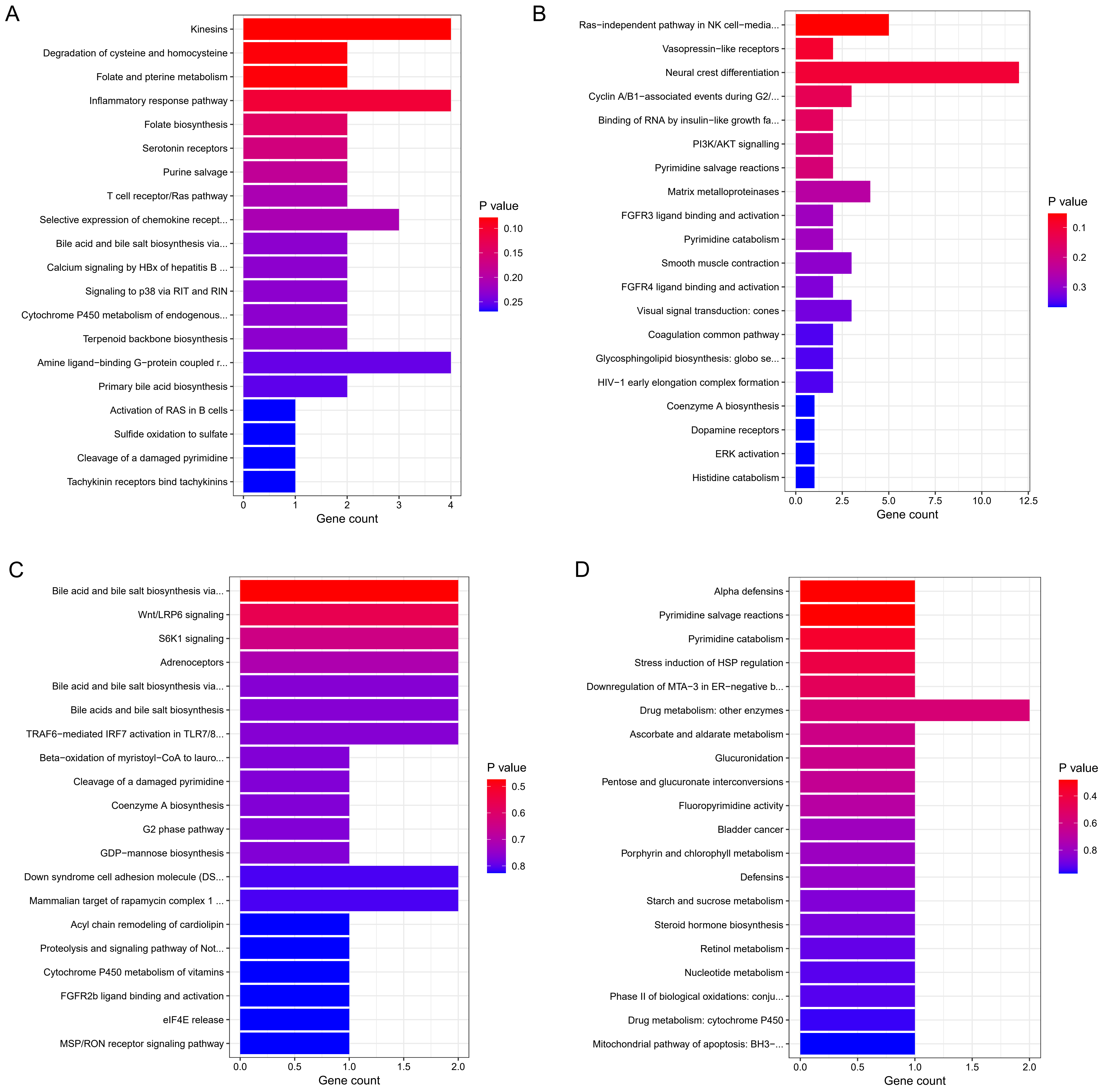

### Supplemental Figure 4

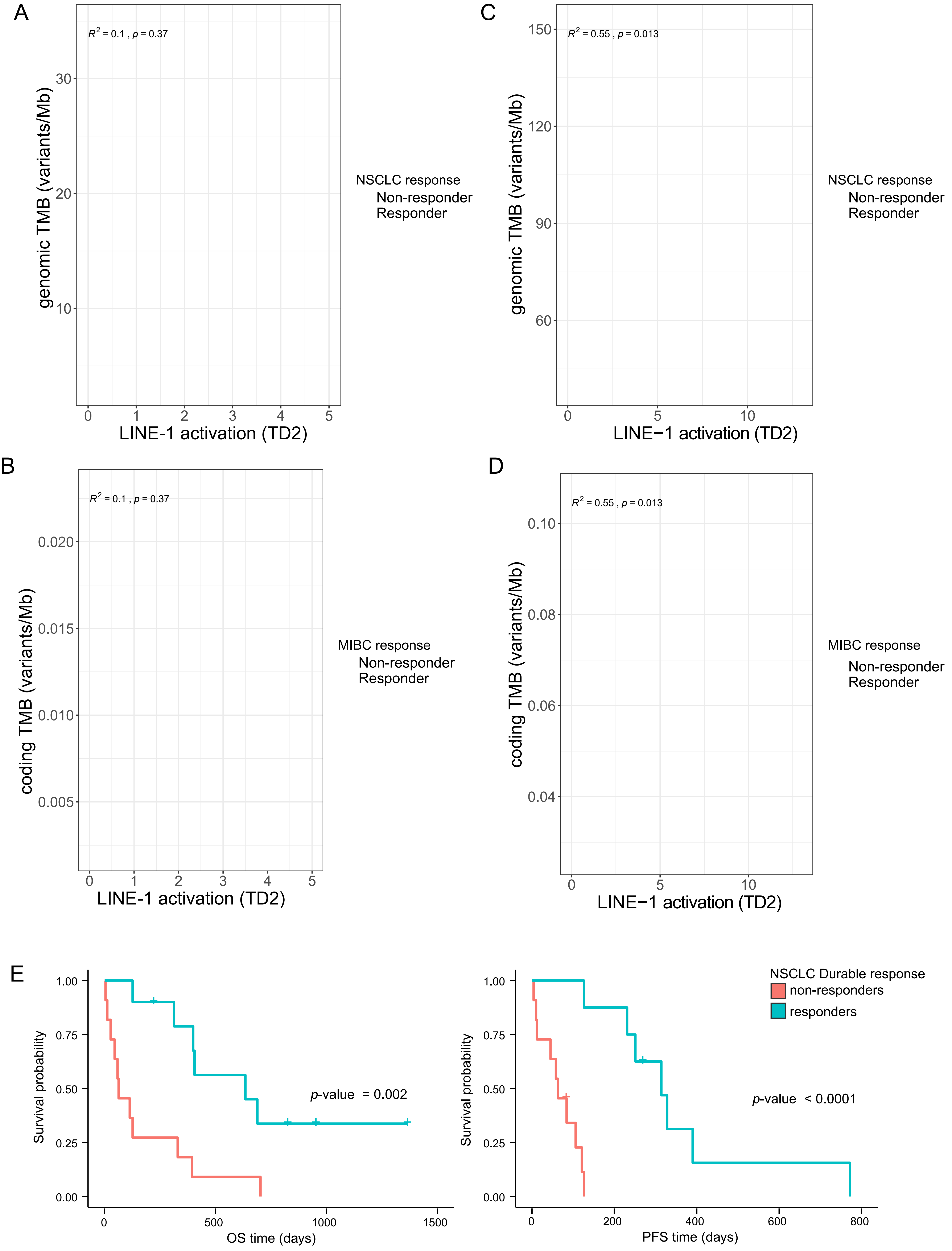
